# Gene Expression Landscapes Driving Early Life Stages of the Keystone Seagrass *Posidonia oceanica*

**DOI:** 10.64898/2026.03.13.711526

**Authors:** Gianmarco Valenti, Alberto Sutera, Emanuela Dattolo, Francesco Carimi, Gabriele Procaccini, Francesco Mercati, Guglielmo Puccio, Roberto De Michele

**Affiliations:** Institute of Biosciences and Bioresources (IBBR), CNR., Via Ugo La Malfa 153, 90146 Palermo, PA, Italy; Department of Integrative Marine Ecology, Stazione Zoologica Anton Dohrn, Villa Comunale, Naples 80121, Italy

**Keywords:** development, gene ontogeny, photosynthesis, *Posidonia oceanica*, seagrass, seed, seedling, transcriptomic, WGCNA

## Abstract

Seagrasses are marine angiosperms forming extensive underwater meadows that provide habitat, stabilize sediments, store carbon, and protect coastlines. *Posidonia oceanica* is the endemic foundation seagrass species of the Mediterranean, yet its meadows are rapidly declining. Despite its ecological importance, the molecular basis of *P. oceanica* development remains poorly understood.

Here, we analyzed gene expression in roots, leaves, and seeds across four developmental stages, revealing strong tissue-specific patterns and temporally regulated expression dynamics. Leaves exhibited active regulation of photosynthesis-related processes, while roots were enriched in pathways linked to carbohydrate metabolism and cell wall biogenesis, supporting primary root growth and establishment. Seeds retained metabolic activity, with glycolytic enzymes indicating readiness for germination. Temporal analyses identified a major transcriptional shift, with distinct gene sets sequentially activated during early establishment and late maturation across tissues.

Weighted Gene Co-expression Network Analysis identified modules strongly associated with specific tissues and developmental transitions, highlighting key hub genes involved in photosynthesis, metabolism, cell wall remodeling, and protein synthesis. Together, these results reveal complex, temporally coordinated regulatory networks underlying *P. oceanica* development

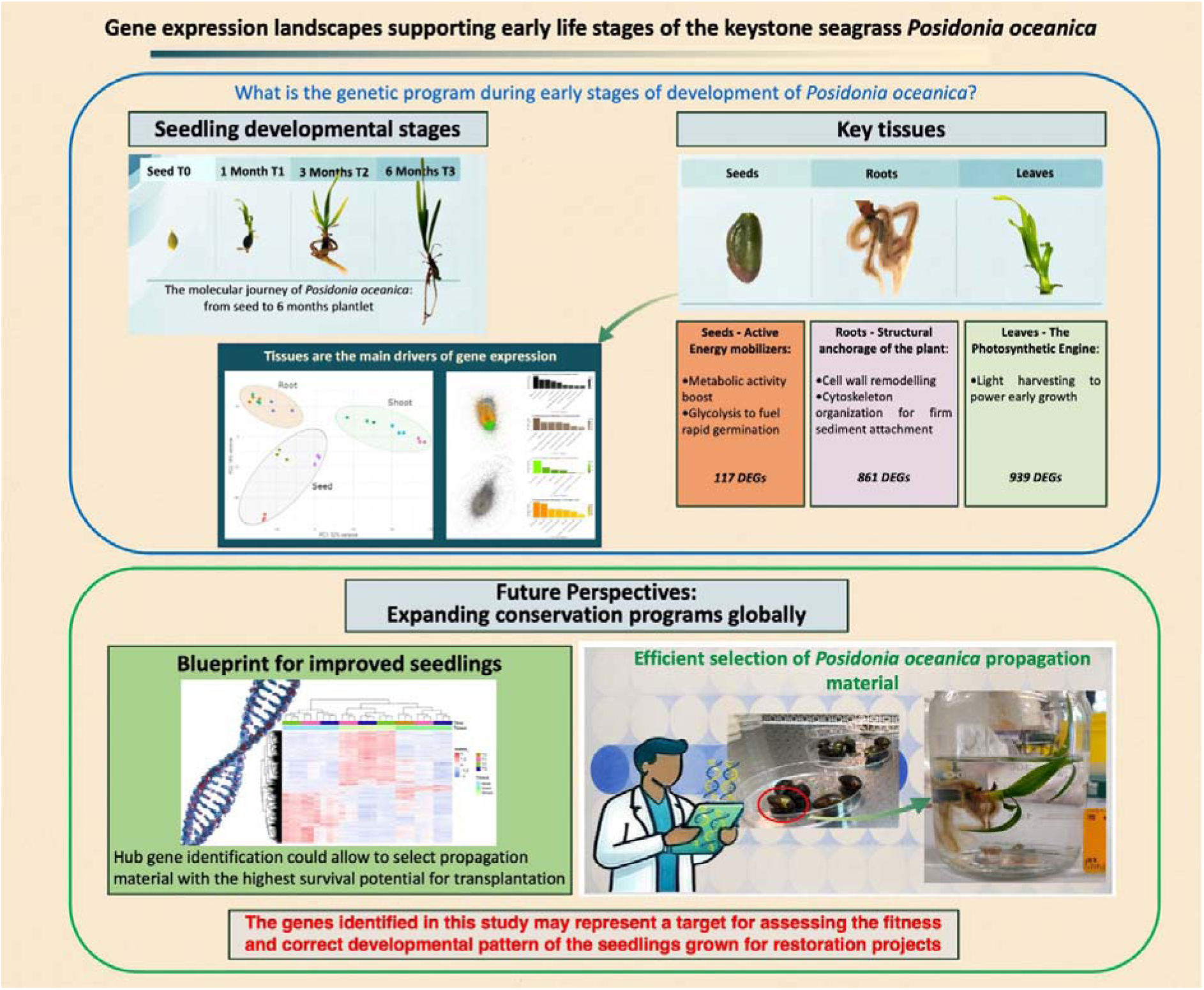

**Highlight:** Gene expression analyses reveal tissue-specific and temporally regulated networks driving *Posidonia oceanica* development, identifying key pathways and hub genes coordinating early establishment and late maturation across roots, leaves, and seeds.

## 1. Introduction

Seagrass meadows are among the most valuable coastal ecosystems on Earth. They act as an erosion barrier that dampens swell, limiting sediment movement on the seabed and actively participating in the sedimentary balance of the beach by supplying biogenic sand(De Falco *et al*., 2017). Meadows provide food and protection to fishes and invertebrates, thus serving as incubator for the fishing industry (Mtwana Nordlund *et al*., 2016). Additionally, they act as carbon sink, mitigating the global increase of carbon dioxide (Duarte and Krause-Jensen, 2017). *Posidonia oceanica* (L.) Delile is the dominant seagrass in the Mediterranean. However, anthropogenic pressure and climate change are menacing meadows health and distribution, with documented regression reaching 34% over the past fifty years (Telesca *et al*., 2015). Targeted actions are therefore essential to mitigate the factors causing regression and to implement conservation practices that may help preserve these habitats’ key coastal ecosystem functions.

*P. oceanica* reproduces both vegetatively, by lateral expansion of its rhizomes, and sexually, by releasing large buoyant fruits, which dehisce and release a single seed. Unlike other seagrass species, *P. oceanica* seeds are not dormant, and seedling development begins already within the fruit (Orth *et al*., 2000; Belzunce *et al*., 2005). Both lateral cuttings and seeds collected on beaches have been used as propagation material for restoration efforts (Boudouresque *et al*., 2021). A seed-based strategy is cost-effective and preserves the genetic diversity of populations. However, it is hampered by a rapid decrease in seed viability in stranded seeds, unless protected by fruits or shaded (Sutera *et al*., 2024). Early seed collection is essential; after that, seed viability can be preserved for several months under cold and lit conditions, allowing an effective decoupling of seed collection from seedling development and transplant (Sutera *et al*., 2025).

During early seedling development, leaves and the primary root emerge from seed, which persists and nourish the plantlet both with the starchy hypocotyl and with active photosynthesis up to the first year (Balestri *et al*., 2009; Celdran, 2013; Celdran *et al*., 2015; Guerrero-Meseguer *et al*., 2018). Leaves rapidly elongate, and new leaf pairs form (Sutera *et al*., 2025). Secondary roots also emerge and grow (Belzunce *et al*., 2008).

Recently, the genome of *P. oceanica* and other seagrass species has been sequenced, revealing processes of genomic adaptation to underwater marine life for plants originally evolved on land (Ma *et al*., 2024*a*). However, so far only few studies have explored the patterns of gene expression in *P. oceanica* in physiological and stress conditions, and they were all performed on adult plants (Dattolo *et al*., 2013; Entrambasaguas *et al*., 2017; Procaccini *et al*., 2017; Marín-Guirao *et al*., 2019; Ruocco *et al*., 2021; Pazzaglia *et al*., 2022, 2025; Santillán-Sarmiento *et al*., 2023).

Therefore, a global transcriptomic picture of the plant’s earliest developmental stages is lacking. This study addresses this critical gap by providing the first transcriptome analysis during seed germination and early seedling development in *P.oceanica* with a tissue-resolved resolution. The identification of key genes and regulatory metabolic pathways that play pivotal roles in the plant’s ontogeny might assist the selection of the best performing seedling as propagation material for meadow restoration projects.

## 2. Materials and methods

### 2.1 Seedling growth and experimental design

Stranded fruits were collected from the shore of Trabia (37.994 N, 13.666 E)) on May 19, 2022. The seeds were stripped from the fruit and placed in controlled conditions in an aquarium (40 L) with artificial seawater (3.8% salinity), biological filtration, controlled temperature (18°C) and growth light (300 lx with a photoperiod of 14 h/10 h light/dark), with no substratum. The first time point analyzed (T0) consisted of three replicates of freshly collected seeds. The second time point (T1) involved the analysis of three biological replicates for each tissue (i.e. leaf, root and seed), selected from seedlings one month after being placed in the aquarium. Two additional time points were analyzed following the same procedure, but with seedlings selected at three months (T2) and six months (T3) after germination. For simplicity, throughout the manuscript we refer to the seed remnant, attached to a developed seedling, as seed. Leaf and root length were measured on pictures by ImageJ software, on five replicates. Within each tissue, length across different developmental stages were compared by Kruskal-Wallis test, followed by Pairwise Mann-Whitney tests (alpha = 0.05).

### 2.2 RNA extraction

Leaf, root and seed samples were collected at each time point, carefully avoiding cross-tissue contaminations, and stored at -80°C pending processing. From each sample, aliquots of approximately 60-80 mg were grinded in liquid nitrogen. The RNA of leaf and root tissues was extracted from ground material with the Aurum™ Total RNA Mini Kit (BIO-RAD), following the manufacturer’s protocol for Plant Tissue. Seed samples contained a large amount of starch, which clogged the purification columns; therefore, the initial extraction phase was performed using the Small Scale RNA Isolation procedure outlined in the datasheet of the PureLink® Plant RNA Reagent (Thermofisher). Subsequently, the aqueous phase was processed using the Aurum™ Total RNA Mini Kit. Both purity and concentration of total RNA were assessed using a BioTek Synergy Plate Reader spectrophotometer. RNA integrity was assessed by electrophoresis on a 1% agarose gel. Due to their poor RNA quality, the samples obtained from seed tissues at T2 were excluded from the analysis, resulting in a final dataset of 27 samples.

### 2.3 RNA-seq analysis

The prepared samples were sequenced using an Illumina NovaSeq 6000 platform, following the Illumina protocol, resulting in the generation of paired-end reads (Biodiversa, Italy). The quality of the raw sequencing data was assessed using FastQC (Wingett and Andrews, 2018). Reads were then trimmed using TRIMMOMATIC (Bolger *et al*., 2014)with default parameters obtaining high quality clean reads with a median per-base sequence quality of 36 (Phred Score) and no adapter sequences. Sequencing revealed a skewed GC content indicative of eubacterial presence. This was expected because the seedlings were grown under non-sterile conditions and because *P. oceanica* seeds already harbor a diverse community of endophytes (Crucitti et al., 2026).In any case, mapping the reads against the reference *P. oceanica* genome ensured the removal of non-plant sequences. Reads were mapped to the newly assembled *P. oceanica* genome (Ma et al., 2024) by using the tool STAR (Dobin *et al*., 2013), with default parameters. Finally, mapped reads were assigned to genomic features using featureCounts (Liao *et al*., 2014), with default parameters.

DESeq2 was utilized to perform the Differential expression analysis (Love *et al*., 2014). The raw counts were first normalized through DESeq2 size factor normalization and transformed using the the variance-stabilized transform (VST). Principal component analysis (PCA) was performed on VST data. Tissue-specific Differentially Expressed Genes (DEGs) were obtained through pairwise comparisons between tissues at each time point, using a false discovery rate (FDR) threshold of 0.001 and a log□ fold change threshold of > 2 for up-regulated DEGs and < -2 for down-regulated ones. The results of these multiple comparisons were visualized and analyzed using an UpSet plot generated with the ComplexHeatmap R package (Gu, 2022)to reveal shared DEGs across conditions. In parallel, to capture genes with significant trends across different time points, a likelihood ratio test (LRT) was employed using a multi-factor design. For this test, the full model (∼ Tissues + Time + Tissues:Time) was tested against a reduced model excluding the interaction term (∼ Tissues + Time).

### 2.4 Functional and co-expression network analysis

Gene Ontology (GO) enrichment analysis was performed using the R package ClusterProfiler (Yu *et al*., 2012) with an adjusted p-value cutoff of 0.01 which was calcuated using the Benjamini-Hochberg method. Results were visualized using the R package ggplot2(Wickham, 2011). Weighted Gene Co-expression Network Analysis (WGCNA) was applied to VST data to identify co-expression modules. A soft-thresholding power of 8 was selected to achieve scale-free topology (Supplementary Figure S1). Modules were identified using dynamic tree cutting with a minimum size of 30 genes. For each module, module-trait associations were calculated as the Pearson correlation coefficient between the module eigengene (the principal component of the module expression profile) and a binary matrix with all the experimental conditions. Network visualization was performed in Cytoscape and hub genes were defined as the top 5% most connected genes within each module.

### 2.4 Statistical analysis

The number of leaves and roots for each developmental stage is presented as mean and standard deviation (n = 5 independent seeds/seedlings). Pairwise differences were statistically tested by t-test. Maximum leaf and root lengths among developmental stages were compared by Mann-Whitney tests, alpha = 0.5.

For the transcriptome analyses, n = 3 independent seeds/seedlings. Statistical processing of transcriptome data has been described in detail in the previous section.

## 3. Results

### 3.1 Seedling development

Seeds germinated rapidly in aquaria. A pair of leaf primordia was already apparent in the seed apex in many seeds once the fruit was removed (Supplementary Figure S2), and rapidly expanded during growth. After one month, six leaves were already present, and an additional pair appeared at later times (Figure 1A). Leaves elongated, reaching about 16 cm after six months (Figure 1B). Primary roots emerged on the opposite side of the seed in most samples, though not in all of them, but its growth stopped already after the first month (Figure S2). Secondary, adventitious roots developed later at the rhizome junction, increasing in number and length along time (Figure 1A, B). At six months, branching of secondary roots was also evident (Figure 1C, Figure S2).

**Figure 1:**
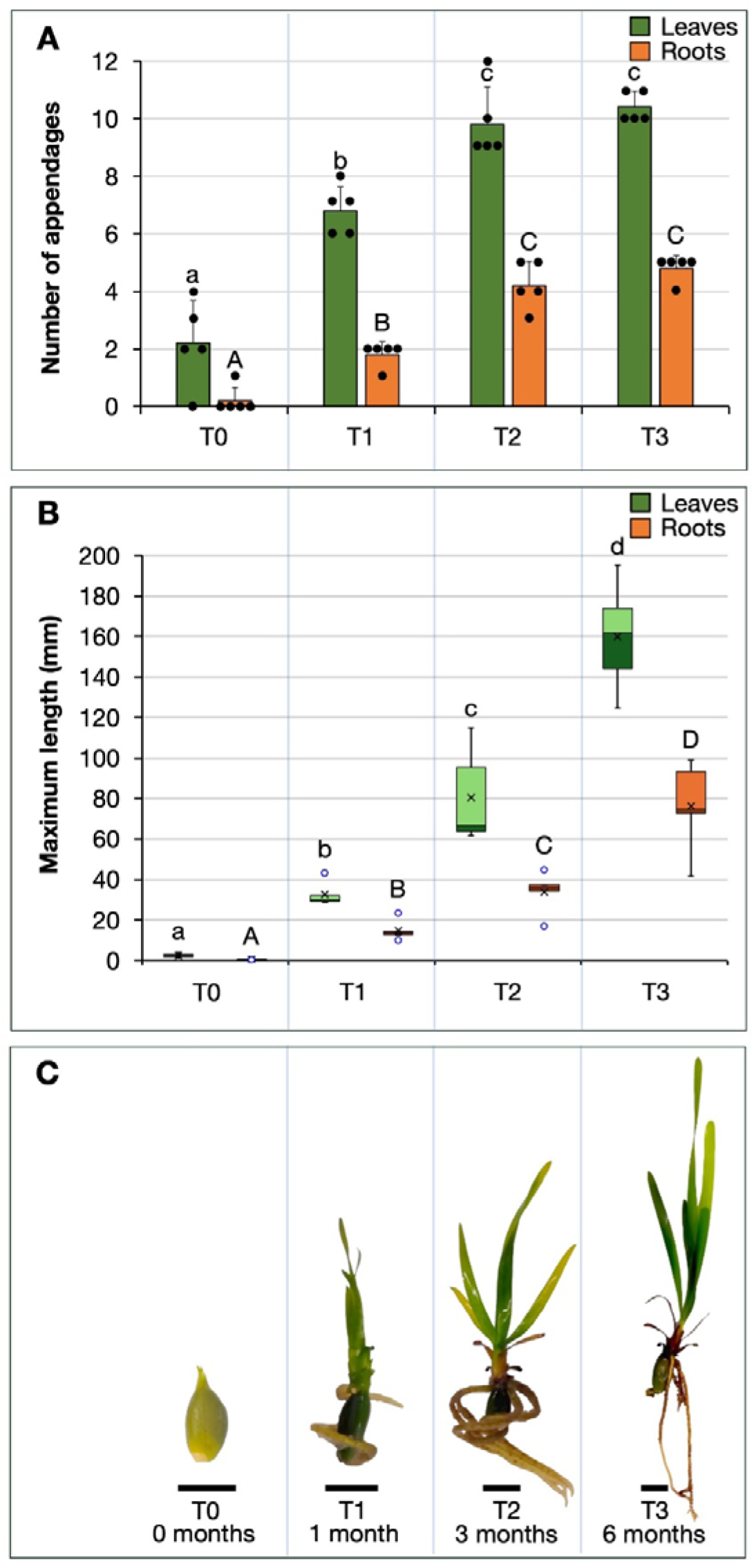
Seedling development. A: Number of leaves and roots; means and standard deviations are presented; dots represent the actual data values. B: maximum leaf and root length; means are presented as crosses, medians as lines dividing boxes. C: representative images of seedling at each developmental stage. Bar = 1 cm, n = 5 biological replicates. Different letters indicate significant differences among time points within each tissue, as assessed by t-test (A), and pairwise Mann-Whitney tests (B), alpha = 0.05.

### 3.2 Transcriptional profiling of *P. oceanica* tissues

A total of 1.1 billion paired-end (PE) reads were mapped to the reference genome, averaging 42 million reads per sample. On average, 18,478 ± 294 (SD) unique genes were detected, representing 79% of the total gene set.

The Principal Component Analysis (PCA) of the 27 samples of *P. oceanica* revealed a clear separation of biological replicates according to tissue type. Samples grouped into three distinct clusters corresponding to Root, Leaf, and Seed (Figure 2A). The first two principal components explained the majority of the variance, with PC1 and PC2 accounting for 53 % and 19 %, respectively.

**Figure 2.**
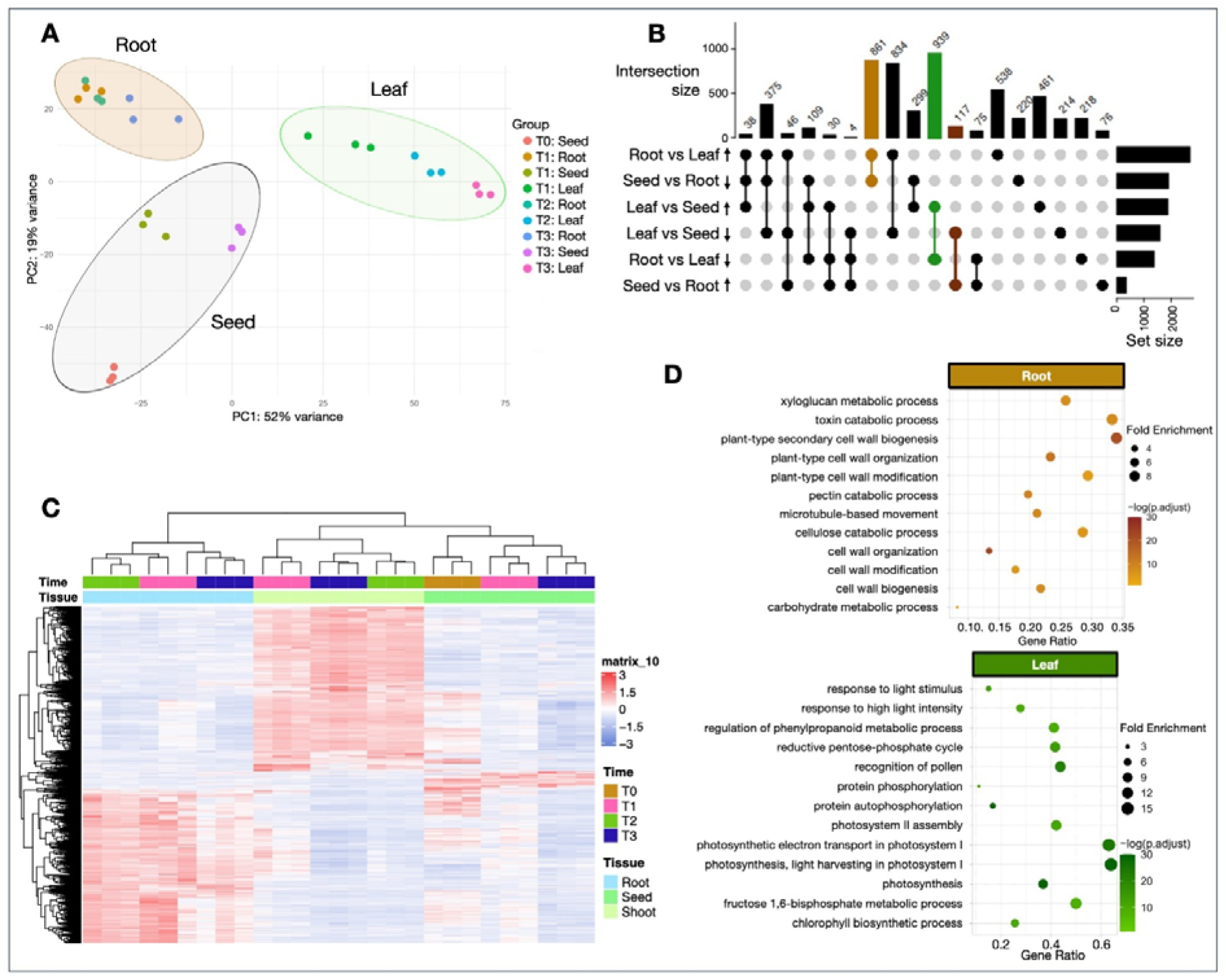
Tissue and temporal distribution of gene expression. A: Principal Component Analysis (PCA) performed on three tissues (root, leaf, seed) at different time points (0; 1; 3; 6 months after germination) of *P. oceanica* seedlings, n = 3 biological replicates. B: UpSet plot displaying the intersection patterns of DEGs identified in pairwise contrasts. Tissue-specific DEGs are color-coded: leaf (green), root (light brown), seed (dark brown). Bars represent intersection sizes, with connected dots indicating contributing comparison sets and numbers on each bar representing the number of DEGs. C: Heatmap showing the expression patterns of tissue-specific genes. D: Dot plot showing GO enrichment results for biological process in leaf and root specific genes.

Pairwise contrast between tissues revealed extensive transcriptional reprogramming during development with a large number of Differentially Expressed Genes (DEGs). The identified DEGs were further filtered in order to differentiate Up regulated genes from down regulated ones (log2 fold change > 2 for upregulated DEGs and < –2 for downregulated ones). Overall, 4,009 (2,660 up-regulated and 1,349 down-regulated) DEGs were detected in the root vs leaf contrast, 2,101 (304 up-regulated and 1,797 down-regulated) in seed vs root, and 3,397 (1,794 up-regulated and 1,503 down-regulated) in leaf vs seed (Supplementary Table S1).

Restricting the analysis to upregulated DEGs in each tissue allowed the identification of distinct tissue-specific gene sets, comprising 939 DEGs unique to leaves, 861 DEGs unique to roots, and 117 DEGs unique to seeds (Figure 2B; Supplementary Table S2). A heatmap of these tissue-specific DEGs displayed their expression profiles across the four developmental stages (T0–T3). Hierarchical clustering revealed consistent grouping of samples by both tissue type and developmental stage (Figure 2C).

To gain insight into the biological roles of these DEGs, Gene Ontology (GO) enrichment analysis was performed. Leaf-specific DEGs were predominantly enriched in photosynthesis-related processes, photorespiration, response to light stimulus and metabolic regulation. Root-specific DEGs were enriched in carbohydrate metabolism and cell wall-related processes, including *xyloglucan metabolic process* and *cell wall organization* (Figure 2D). Seed-specific DEGs showed a more restricted enrichment, with only one significant term detected in biological processes, *polysaccharide catabolic process* (Supplementary Table S2).

### 3.3 Time-dependent modulation of gene expression during tissue development

A Likelihood Ratio Test (LRT) was applied to identify genes showing tissue- and time-specific transcriptional dynamics, revealing 4,642 DEGs (padj < 0.001). Hierarchical clustering grouped these genes into nine clusters: Cluster 1 (382 DEGs), Cluster 2 (1,080), Cluster 3 (365), Cluster 4 (832), Cluster 5 (199), Cluster 6 (210), Cluster 7 (588), Cluster 8 (840), and Cluster 9 (146) (Supplementary Table S3).

Analysis of expression patterns (Figure 3A) highlighted tissue- and time-dependent trends. Clusters 1, 3, and 9 showed progressive upregulation in leaves and downregulation in roots and seeds. Cluster 2 remained stable in roots and seeds but was downregulated in leaves. Clusters 4, 5, and 7 exhibited overall downregulation over time, with moderate increases in roots but pronounced decreases in leaves and seeds.

**Figure 3.**
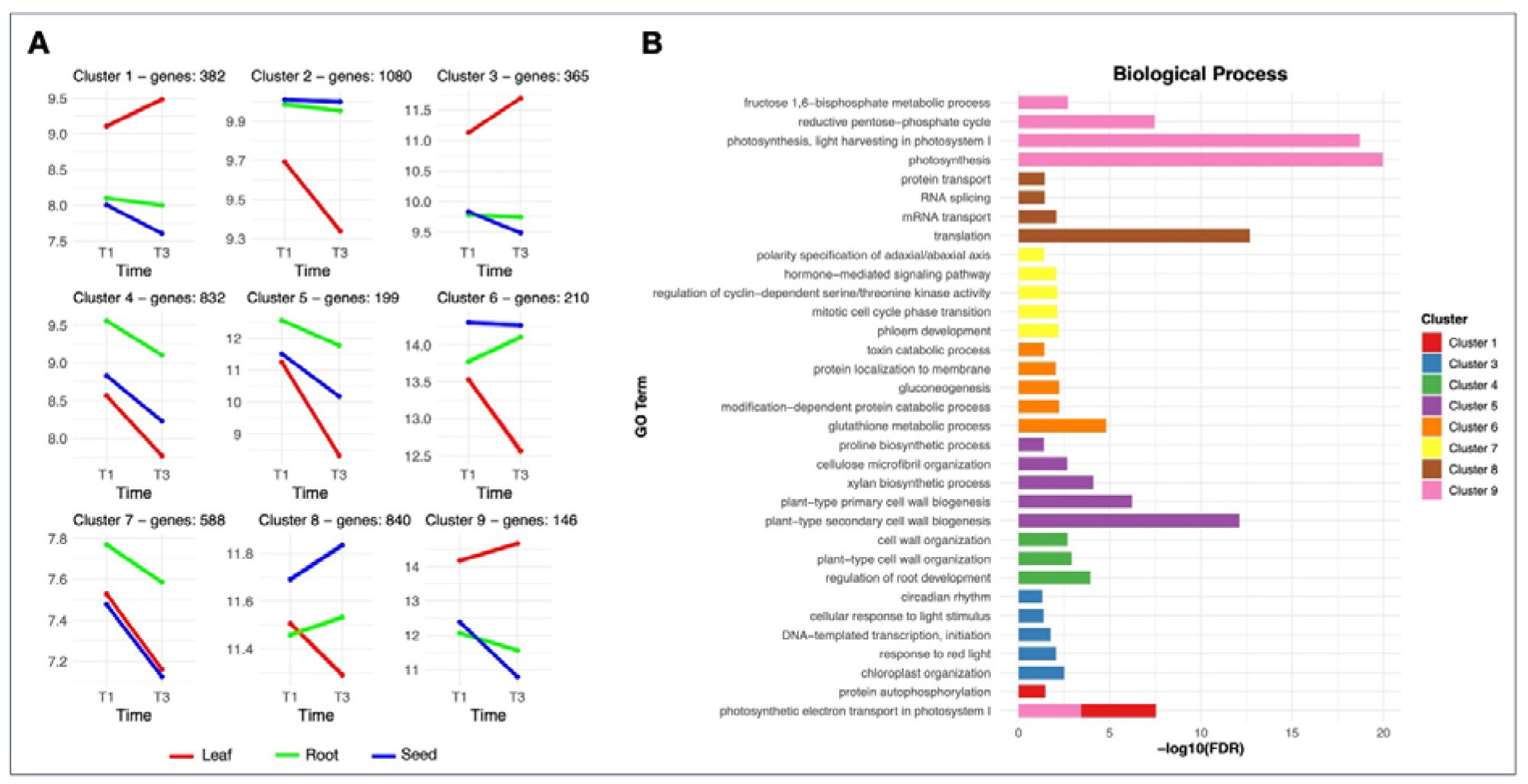
Expression patterns and associated GO terms. A: Dotplot showing temporal trends of clusters obtained by LRT analysis for the interaction between Time and Tissue. B: Barplot showing the 5 most enriched GO terms (BP) of identified clusters.

Functional enrichment of these clusters (padj ≤ 0.01) revealed biologically relevant patterns (Figure 3B), with different temporal regulation in the different tissues. In leaf, clusters upregulated over time were enriched in photosynthesis-related terms, including photosynthetic electron transport in photosystem I and cellular response to light stimulus. Other clusters were downregulated over time, such as Clusters 4 and 5, associated with cell wall development and growth (cell wall organization, primary/secondary cell wall biogenesis), and Cluster 7, involved in hormonal regulation of cell division (hormone-mediated signaling, cyclin-dependent kinase regulation). Cluster 6, enriched in metabolic and stress response processes (gluconeogenesis, glutathione metabolic process) showed contrasting regulation between root and leaf, up- and down-regulated, while in seeds it was consistently up-regulated. Finally, Cluster 8, predominantly linked to gene expression regulation (RNA splicing, mRNA transport) showed a marked up-regulation particularly in seed, with a similar trend also observed in root, while in leaf a downregulated profile was recorded.

### 3.4 Co-expression network analysis

To uncover co-expression gene networks underlying *P. oceanica* development, we performed a Weighted Gene Co-expression Network Analysis (WGCNA). The whole transcriptimic profiles, characterized by a total of 18,299 genes, were grouped through a clustering analysis, and a sample-correlation matrix was calculated. The analysis led to the identification of a total of 11 co-expression modules after the merging phase, each represented by a distinctive color, as shown in Figure 4. These ranged from 91 (module ivory) to 5189 genes (module Bisque4), with the second largest modules being orange (4,531 genes).

**Figure 4.**
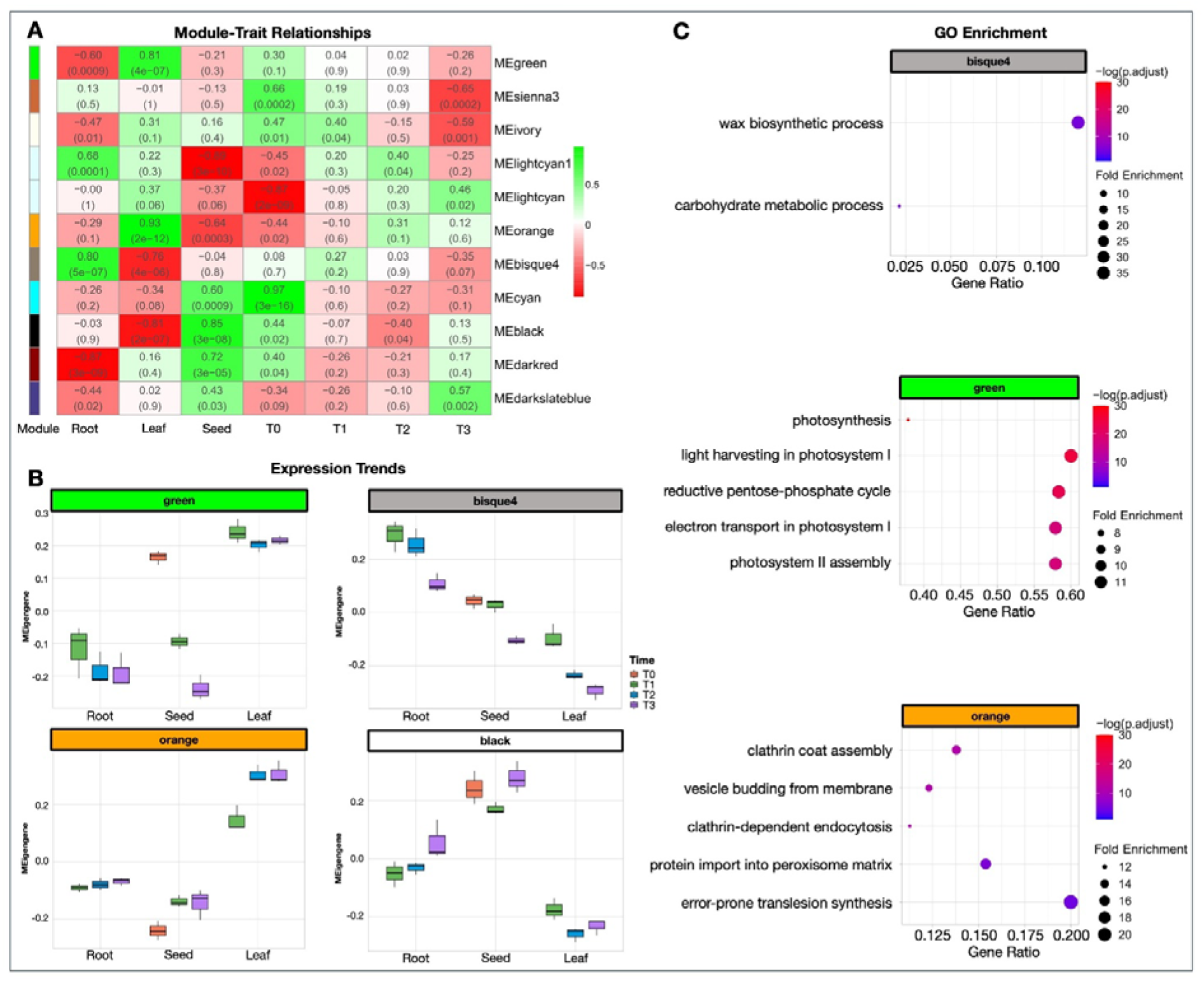
A: Weighted Gene Co-expression Network Analysis of P. oceanica samples. Module-trait relationships revealed by the Pearson correlation coefficient. The leftmost color column indicates different co-expression modules. The numbers in the figure indicate the correlation between the modules and traits, and the numbers in the parentheses are the correlation P-value. B: Module eigengene temporal trend across WGCNA modules in which tissue-specific genes are grouped (green, orange, black and bisque4). C: Biological Process GO enrichment analysis of the corresponding modules through dot-plot showing GO enrichment analysis of modules bisque4, green and orange.

The correlation analysis between modules eigengenes and the experimental conditions (as binary matrix) comprising time (T0, T1, T2, T3) and tissue (root, leaf, seed), allowed to select specific modules correlated to time points and tissues. Four modules showed significant correlations with specific tissues (|r| > 0.6): bisque4 and lightcyan1 with roots, orange and green with leaves, and four smaller modules (darkslateblue, darkred, black, and cyan) with seeds. Modules showing significant correlations with developmental time were identified. The early stage (T0) featured sienna3, ivory, cyan, black, and darkred modules, while ivory was uniquely associated with the T1 time point. Subsequent stages displayed stage-specificity, with lightcyan1 dominating at T2 and coordinated activity between darkslateblue and lightcyan emerging during late development (T3).

To identify modules with tissue-specific expression patterns, DEGs from pairwise contrasts were mapped onto WGCNA modules. Almost 96% of the root specific DEGs were assigned to the bisque4 module (823 DEGs) which was enriched in functions involved in “carbohydrate metabolic process” and “wax biosynthetic process”. Leaf-specific DEGs (930) were distributed across the green and orange (∼ 33% and 55%, respectively) modules. The green module was enriched in GO terms related to photosynthetic activity, such as “photosynthesis”, “light harvesting in photosystem I” and “photosystem II assembly; the orange module for detoxification mechanisms, including “protein import into peroxisome matrix” and vesicle-mediated endocytosis/exocytosis processes, such as “clathrin-dependent endocytosis”.

Seed-specific DEGs (101) were primarily mapped to the black module (58 DEGs) which did not show significant enrichments.

### 3.5 Co-expression modules involved in tissue differentiation and identification of key molecular regulators

Co-expression modules were further characterized through correlation analysis of module eigengenes, network analysis, and functional analysis of hub-genes (Figure 5; Supplementary Figure S3, Supplementary Table S4). The eigengene correlation analysis reveals the global transcriptional architecture of *P. oceanica* development, with modules showing similar expression profiles clustering together (Figure 5A). The hierarchical clustering of module eigengenes reveals a sharp architectural divergence between tissue-specific transcriptional programs. Specifically, seed-and root-associated modules (black and bisque4) are clustered independently from leaf-associated modules (green and orange). While the latter two show a high correlation coefficient (r = 0.62), their separation into distinct sub-clades highlights their specialized biological roles: photosynthesis for the green module and broader metabolic signaling for the orange module.

**Figure 5.**
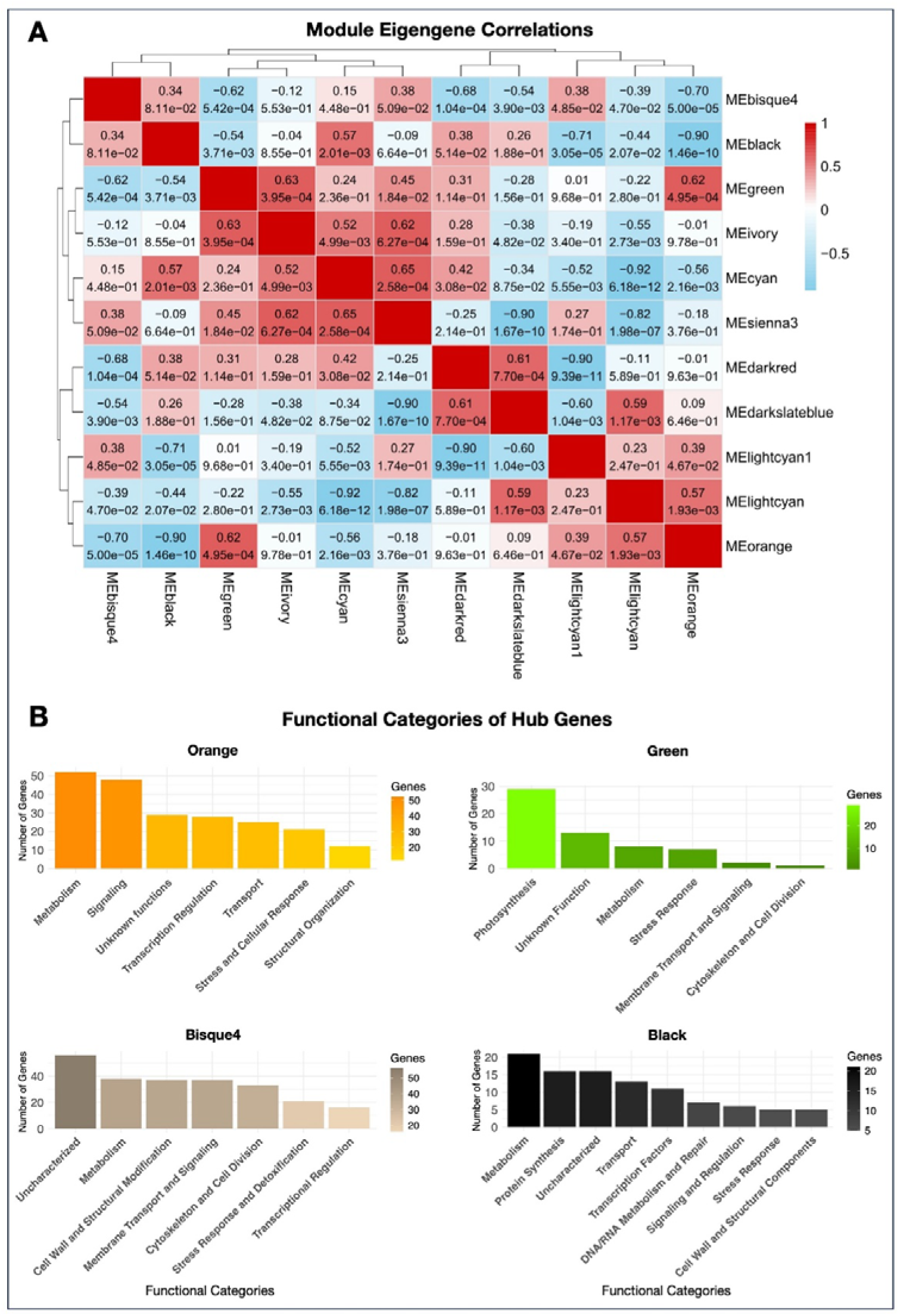
Correlation and functional analyses of hub genes. A: Heatmap of Module–Module Eigengene Correlations. The dendrogram and color scale represent the hierarchical relationship between Co-expression Network Analysis modules, where red indicates positive and blue indicates negative eigengene correlation. B: Barplot showing the function of hub genes identified in the black, bisque4, green, orange co-expression modules of *P. oceanica* through Weighted Gene Co-expression Network Analysis (WGCNA). The x-axis reports the functional categories of the identified hub genes, while the y-axis indicates the corresponding number of genes for each category.

The seed development-associated black module was dominated by transcriptional control (11 genes), including regulators such as *ZAT9*, which orchestrate embryo development through gibberellin and ethylene responses (Joseph *et al*., 2014). Protein turnover specialists (16 genes) maintained proteostasis, while DNA/RNA processors (7 genes) ensured genetic stability. Metabolic hubs (21 genes) fueled growth via glycolysis and the pentose phosphate pathways, supported by transporters (13 genes) for reserve accumulation and stress protectants (5 genes) to shield the developing embryos.

Analysis of the root development-associated bisque4 module highlighted specialized networks for marine adaptation, characterized by structural genes (37 genes) for cell wall remodeling, metabolism regulators (38 genes), and signaling hubs (37 genes) coordinating nutrient uptake. Cytoskeletal organizers (33 genes) enabled cell division, while stress responders (21 genes) mitigated marine oxidative stress and transcriptional regulators (16 genes) guided root patterning.

The leaf development-associated modules showed distinct specializations. The green module concentrated on chloroplast function (29 genes) and metabolic support (8 genes), with additional roles in stress resilience. By contrast, the orange module exhibited broader regulatory roles, primarily in metabolic diversification (49 genes), signaling networks (48 genes), and transcriptional control for processes such as iron homeostasis and morphogenesis.

## 4. Discussion

Our study was aimed to untangle the molecular processes underlying the development of *P. oceanica* seedlings across three key tissues (root, leaf, and seed) and four developmental stages, by combining differential gene expression analysis, temporal dynamics modeling via a Likelihood Ratio Test (LRT), and Weighted Gene Co-expression Network Analysis (WGCNA). The integrated transcriptomic approach revealed a clear tissue-specific pattern in gene expression, with distinct transcriptomic modules dedicated to photosynthesis in leaves, structural and metabolic adaptation in roots, and energy mobilization in seeds, demonstrating how specialized functions shape different tissues, alongside dynamic temporal regulation across developmental stages.

The analysis demonstrates that tissue type is the primary determinant of transcriptional variation, accounting for the majority of the variance observed in the Principal Component Analysis, where the clear separation of samples by tissue type, which accounted for the majority of the variance (72%), unequivocally identifies tissue identity as the primary driver of transcriptional variation in developing *P. oceanica*. This finding is consistent with transcriptomic studies in model plants like *Arabidopsis thaliana* and crop species, where tissue origin consistently emerges as the strongest driver of gene expression patterns (Brady et al., 2007; Schmid et al., 2005).

Notably, the PCA revealed a clear pattern in which the T0 time point marks the starting phase of seedling differentiation into leaf and root. Unlike seeds at later stages, T0 seeds contain leaf and root primordia, and accordingly they occupy an intermediate position between fully developed leaf and root tissues.

Time-tissue interactions further clarified functional shifts: clusters 1, 3, and 9, enriched for photosystem I/II genes, highlighting the contribution of leaf tissues to photosynthesis, consistent with molecular mechanisms underlying photosynthetic acclimation to light availability as described in *P. oceanica* (Procaccini *et al*., 2017). Conversely, clusters 4, 5, and 7, associated with cell growth and hormonal regulation, reflected root developmental processes, corroborating brassinosteroid-mediated salinity responses (Rodríguez-Rojas *et al*., 2024)

### 4.1 Sequential Activation of Transcriptional Programs During Development

The enriched functions of differentially expressed genes identified by Likelihood Ratio Test comprise clusters of GO terms highly linked to the functional specialization of the individual tissues analyzed during plant development.

Clusters 1, 3, and 9, showing progressive upregulation in leaf tissues, are enriched in terms related to photosynthesis, including electron transport in photosystem I and cellular response to light stimulus. This result confirms the central role of leaves during the early development of *P. oceanica* and the importance of regulating photosynthetic function during development, consistent with findings in other angiosperms such as pearl millet (B. Wu et al., 2021),where a significant enrichment of genes linked to photosynthetic activity was observed.

In contrast, cluster 4, 5, and 7, associated with cell wall development, show a decreasing up-regulation trend in all tissues, with higher expression levels in the root. This suggests that the regulation of cell wall production is important for the development of all tissues, but particularly for the development of the root system. This trend is analogous to that seen in other plants like *Maize* (Somssich *et al*., 2016; Stelpflug *et al*., 2016). This pattern likely corresponds to the end of the initial cell elongation phase, a hormonally regulated process that finds support in cluster 7, underscoring the importance of hormonal regulation during root system development, consistent with findings during root elongation (Ma *et al*., 2024*b*).

Furthermore, the activation in seeds (cluster 6) of key metabolic pathways such as gluconeogenesis and glutathione metabolic process is biologically coherent with the germination phase. Gluconeogenesis is indeed crucial for converting seed lipid reserve into glucose, which is necessary for energy sustenance during early growth (Walker *et al*., 2021), while increased gluthatione metabolism might be related to the regulation of seed germination processes (Koramutla *et al*., 2021).

### 4.2 Metabolic and Signaling Responses Govern Leaf Development

We observed a significant enrichment of genes associated with photosynthetic processes, with 930 genes directly involved in pathways such as “photosynthesis,” “photosystem assembly,” and “photosynthetic electron transport.” This result confirms the high photosinthetic activity developed in *P. oceanica* leaves, as for all higher plants. Consistently, terms related to pigment biogenesis such as “chlorophyll biosynthetic process” and “carotenoid biosynthetic process” are enriched, indicating strong activity in chloroplast assembly and synthesis of photosynthetic pigments, which are essential for early development and sustenance of the developing photosynthetic apparatus. Previous studies have already emphasized the importance of these pathways in leaves during light acclimation, as shown by (Dattolo et al., 2017),while (Ruocco *et al*., 2019)highlighted a central role of carotenoids in photoprotection mechanisms during thermal stress conditions.

The results showed that the photosynthetic tissue (i.e. leaves) actively regulates processes such as the fructose-6-phosphate metabolic process, gluconeogenesis, and the reductive pentose-phosphate cycle to support growth of the entire plant. Similar regulatory pattern have been observed in other angiosperms. For instance, studies on tomato have shown how fructose-6-phosphate metabolism and gluconeogenesis are central to stem development (Cai *et al*., 2016), while in *Arabidopsis,* the reductive pentose-phosphate cycle is crucial for the metabolic balance of leaves (Borghi *et al*., 2019). Moreover, the enrichment of genes associated with “protein phosphorylation” and “protein autophosphorylation” indicates a central role of regulation through kinases and signaling pathways, already observed in response to light acclimation in adult *P. oceanica* leaves (Dattolo *et al*., 2017). Additionally, the involvement of the “phenylpropanoid metabolic process” is relevant because phenylpropanoids contribute to lignification, structural stability, and antioxidant defense of tissues, as described by(Xing *et al*., 2024). Finally, some defensive response terms, such as “response to chitin,” highlight the activation of immune pathways linked to the recognition of fungal elicitors, in agreement with observations by (Ruocco *et al*., 2019).

Further analysis of leaf-associated co-expression modules strengthened these observation. The strong positive correlation observed in the module-eigengene correlation analysis suggests that these modules are part of a synchronized leaf-specific transcriptional branch. This coordination likely ensures that photosynthetic output (green) is tightly coupled with the signaling networks and nutrient homeostasis (orange) required for optimal leaf performance in the marine water column. The green module, specifically linked to leaf tissue, was largely composed of genes involved in photosynthesis (particularly components of photosystems I and II) and other key metabolic processes, in line with previously described depth-related modulation of photosynthetic pathways in *P.oceanica* (Procaccini *et al*., 2017). Key players included ferredoxin, chlorophyll a/b binding proteins, and RuBisCO, which stabilize photosystem II and optimize light responses. Ribosomal genes, such as 50S ribosomal protein L1, further underscored the importance of chloroplast maintenance, aligning with (Tourasse *et al*., 2013). Notably, the absence of transcription factors in this module suggests a streamlined network dedicated to photosynthetic efficiency. The orange module, also associated with leaf tissue, seemed to complement the green module by focusing on leaf transcriptional regulation, exhibiting broader regulatory roles and encompassing transcriptional control, signaling, and metabolic diversification. Enzymes like invertase and cystathionine gamma-synthase 1 mirrored those active in *Z. marina* during vegetative growth (Zhao *et al*., 2025), while stress-responsive transcription factors (AP2/ERF, WRKY, bHLH) and signaling genes (e.g., LRR-RK) reflected adaptations critical for marine persistence, as seen in *Z. marina* (Zhao *et al*., 2025).

### 4.3 Transcriptomic Regulation Underpins Root Structural Development

Our study identified 861 specific DEGs involved in key functions in the development of the root system in *P. oceanica*, such as polysaccharide metabolism, cell wall biogenesis, structural remodeling, and environmental response. The upregulation of genes related to functions such as “xyloglucan metabolic process,” “pectin catabolic process,” and “cellulose catabolic process” are fundamental for producing components that drive the expansion and elongation of the root system. These remodeling processes are consistent with the dynamic nature of the cell wall described for seagrasses, which exhibit specific adaptations in cellulose, hemicellulose, and pectin composition(Pfeifer and Classen, 2020). Such processes are sustained by energy generated through carbohydrate metabolism, which exhibits pronounced seasonal variability and metabolic plasticity in *P. oceanica* (Ismael *et al*., 2023). The high expression of genes involved in cell wall construction and modification, such as “cell wall biogenesis” and “cell wall organization,” suggests that *P. oceanica* roots are engaged in intense deposition of cellulose, lignin, and other structural components, confirming what is known about the composition of cell wall in *P. oceanica* (Pfeifer and Classen, 2020). Finally, consistent with a developing and elongating tissue like the root system, functions related to the cytoskeleton, such as “microtubule-based movement,” highlight the cytoskeleton’s role in regulating cell elongation and orienting the deposition of cellulose microfibrils. These processes provide rigidity and mechanical strength, essential characteristics for ensuring plant stability in the sediment and facilitating nutrient transport. Similar dynamics have been documented in terrestrial plants, where cortical microtubules guide cellulose synthase complexes, ultimately shaping cell wall organization and mechanical strength (Lei *et al*., 2014). The bisque4 module, highly upregulated in roots, further linked wax biosynthesis and carbohydrate metabolism to structural reinforcement, in agreement with findings on lignin-mediated hydrophobicity in *P. oceanica* roots (del Río *et al*., 2022). Hub genes additionally emphasized sterol biosynthesis pathways (e.g., *ATP-citrate synthase,* δ*(24)-sterol reductase*), consistent with light-responsive metabolism described in *Zostera (Kumar et al., 2017)*. The presence of MYB transcription factors resembling those active in salt-stressed *Z. marina* (Zhao *et al*., 2025), suggests that conserved regulatory mechanisms may underlie root specialization across different seagrass species. Interestingly, the eigengene correlation heatmap revealed a distinct transcriptional distance between the root-associated bisque4 module and the leaf-associated modules. This separation highlights the high degree of functional specialization in roots, which operate under a regulatory regime distinct from the autotrophic tissues.

These pathways are fundamental for modulating the elasticity and plasticity of the cell wall during root growth. Interestingly, *P. oceanica* seedling roots have been shown to adhere firmly to the substrate, withstanding force up to 2.4 N, effectively resisting to strong storms(Zenone *et al*., 2020). The adhesion is ascribed to mechanical interlocking of roots, due to the plasticity of root hairs which grow along the microroughness of the substrate, adapting to the micro-topography of the surface, rather than through the secretion of adhesion compounds. This scenario is consistent with our findings, which highlight an intense activity of cell wall remodeling and deposition of microfibrils. Taken together, these results reflect the high functional specialization of the root tissue, not only as an organ for anchorage and absorption but also as a metabolically active district capable of adapting to the often extreme conditions of marine sediment.

### 4.4 Transcriptomic Activity Sustains Metabolic and Regulatory Responses in Seeds

This study enabled the identification of key mechanisms that guide early germination and transcriptomic activity in seeds during development. The seed of *P. oceanica* is an active tissue that maintains its activity at advanced developmental stages. Our results indicate that the breakdown and utilization of starch and sugars provide energy and carbon sources during the early stages of germination. This was evidenced by the analysis of 101 seed-specific differentially expressed genes (DEGs) identified through pairwise comparisons, which are enriched in “carbohydrate metabolism” a process crucial for early development These results agree with observation in the seagrass *Z. marina* (Zhang *et al*., 2023), where carbohydrate-related pathways were markedly during early germination. The black module, though not significantly enriched functionally, harbored genes pivotal for germination, such as carbohydrate-metabolizing enzymes (*enolase*, *pyruvate kinase*). These align with *Z. marina* studies (Zhu *et al*., 2023)highlighting the key role of carbohydrates in energy reserves during early growth. Notably, our transcriptome data reveal sustained metabolic activity in *P. oceanica* seeds even post-germination, suggesting continued resource mobilization, a trait critical for seedling establishment in marine environments. This is confirmed by previous analysis of *P. oceanica* seed content, which report a decrease in carbon, nitrogen and phophorous in seeds in the first months after germination, due to nutrient reallocation to leaves and roots(Balestri *et al*., 2009).

Interestingly, we did not observe modulation of genes involved in photosynthesis in the seed tissue. This is in contrast with multiple physiological observations that indicate seeds as functionally active photosynthetic organs in developing *P. oceanica* seedlings (Celdran, 2013; Celdran *et al*., 2015; Guerrero-Meseguer *et al*., 2018). It is conceivable that photosynthesis is restricted to the outermost cell layer in seeds, and therefore the upregulation of photosynthetic machinery remains unnoticed due to the dilution with the bulk of the inner seed, which is more active in reserve mobilization. The hierarchical proximity between the seed and root modules highlights a shared molecular signature that distinguishes these tissues from the photosynthetic organs. The correlation between the black and bisque4 modules points to their combined role in breaking down external nutrients and reorganizing internal cellular components, which sustains *P. oceanica* development before the photosynthetic apparatus reaches full maturity.

Additionally, the presence of transcription factors like *NAC domain-containing protein 87* and MYB73, analogous to those governing dormancy in seashore paspalum (Wu *et al*., 2024) and fatty acid production in *Arabidopsis*(Yang *et al*., 2025), respectively, further implied dual roles in stress response and developmental control beyond the germination phase. These elements further highlight the complexity and fine-tuned control that govern *P. oceanica* seed activation, positioning the seed as a metabolically versatile organ capable of supporting seedling establishment in a challenging marine environment.

### 4.5 Conclusions

This study focused on transcriptional analyses of seedling ontogeny of the ecologically important seagrass *P. oceanica*. By examining gene expression across roots, leaves, and seeds, we identified distinct tissue-specific signatures and regulatory modules that underpin early development. Our analysis revealed temporally regulated gene clusters governing early and late developmental stages, as well as functionally specialized clusters with dynamic expression over time, illustrating the complex regulatory networks guiding *P. oceanica* development in controlled conditions. Collectively, these results enhance our understanding of the molecular basis of *P. oceanica* development, offering insights into its ecological roles in coastal ecosystems, including sediment stabilization, carbon storage, and biodiversity support. The tissue-specific and temporally regulated gene networks identified here provide a foundation for future functional studies, particularly in elucidating how *P. oceanica* adapts to marine environment. Future research should focus on validating these hub genes through functional genomics and exploring their responses under natural environmental conditions to further inform conservation strategies for this critical Mediterranean species. Furthermore, the genes identified in this study may represent a target for assessing the fitness and correct developmental pattern of the seedlings used in restoration initiatives. Early stages are a bottleneck in viability of *P. oceanica* viability, and meadow restoration is a costly effort which would benefit from the employment of the best performing propagation material.

## Author contributions

RDM: conceptualization; GPu, RDM: data curation; GV, GPu, RDM: formal analysis; FC, GPr, RDM: funding acquisition; AS, RDM: investigation; ED, GPu: methodology; FC, GPr, RDM: project administration; AS, RDM: resources; GPu, RDM: supervision; GV, RDM: validation; GV, FC, GP, RDM: visualization; GV, RDM: writing-original draft; all authors: writing-review and editing.

## Data availability

The RNA-Seq data underlying this article are available in NCBI-SRA, and can be accessed with the reference number PRJNA1434360.

## Supplementary data

**Supplementary Figure S1.**
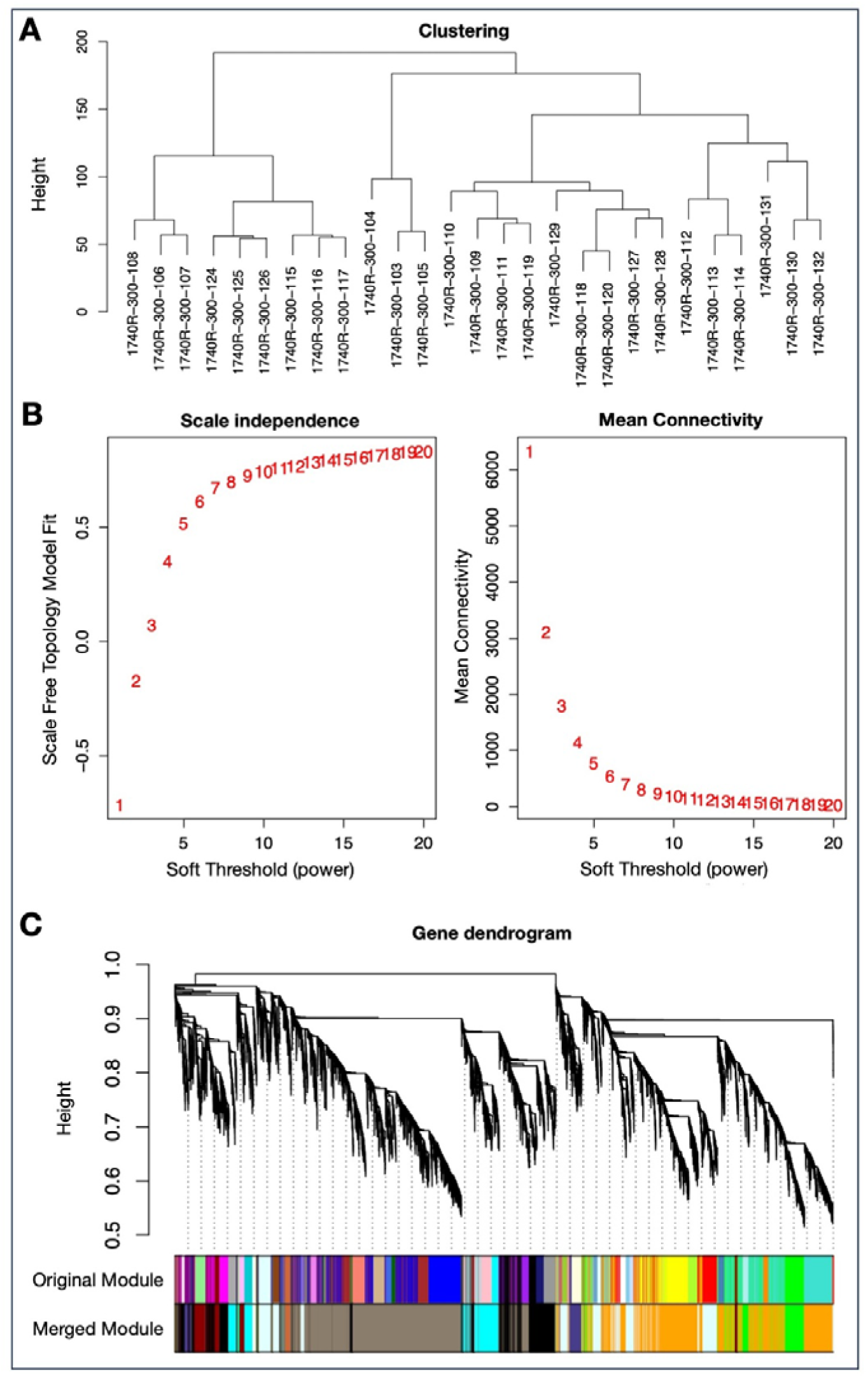
A: Sample clustering based on global gene expression profiles used to assess sample similarity and identify potential outliers prior to network construction. B: Soft-tresholding power selection for WGCNA. The left panel shows the scale-free topology fit index across increasing powers, while the right panel shows the corresponding mean connectivity values. C: Gene dendrogram generated from hierarchical clustering of the topological overlap matrix, with modules assigned through dynamic tree cutting and subsequently merged based on eigengene similarity. Colors denote distinct WGCNA modules.

**Supplementary Figure S2:**
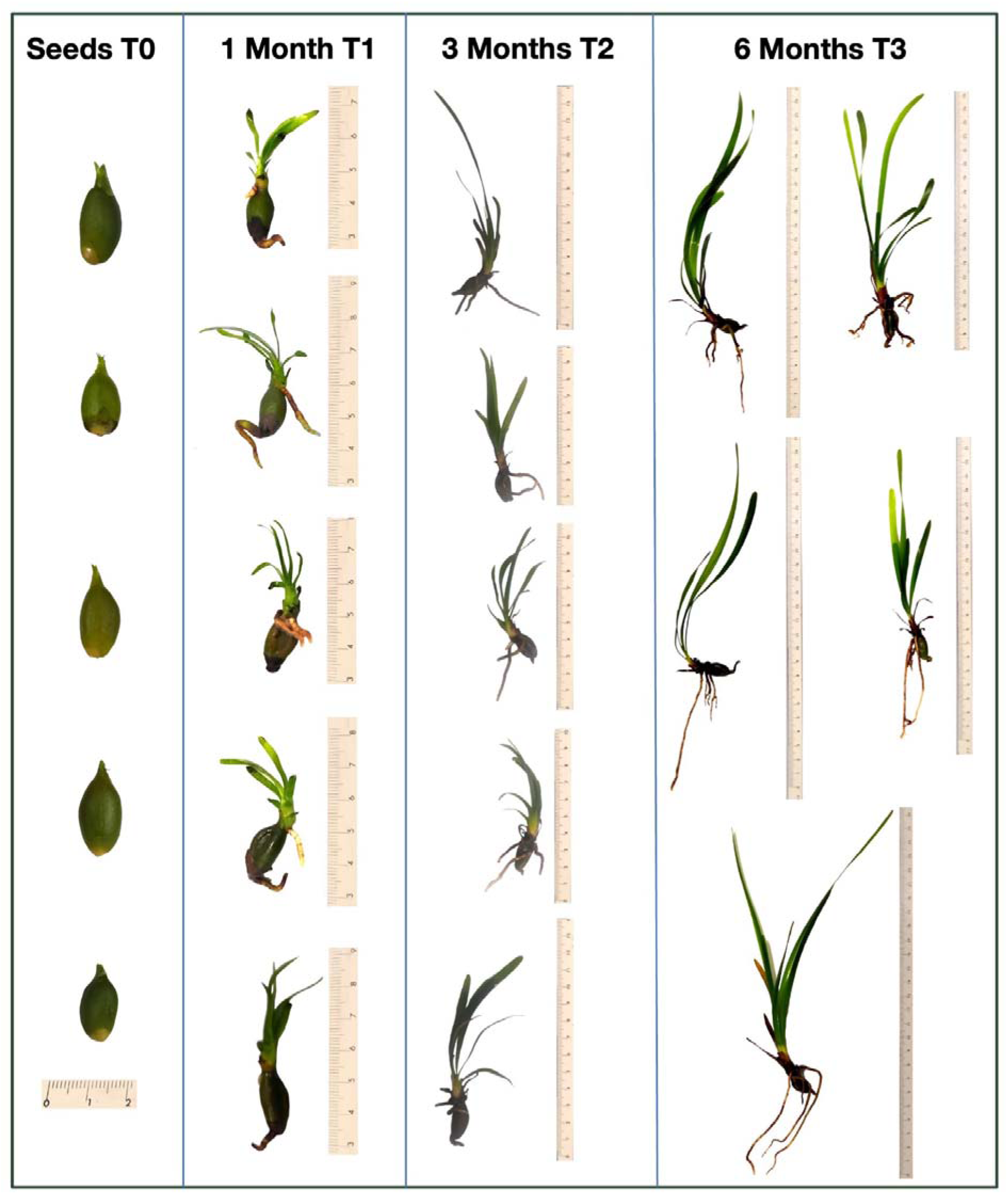
*P. oceanica* seedlings analysed in this study. The three uppermost seedlings of each developmental stage were further processed for the transcriptomic analyses.

**Supplementary Figure S3:**
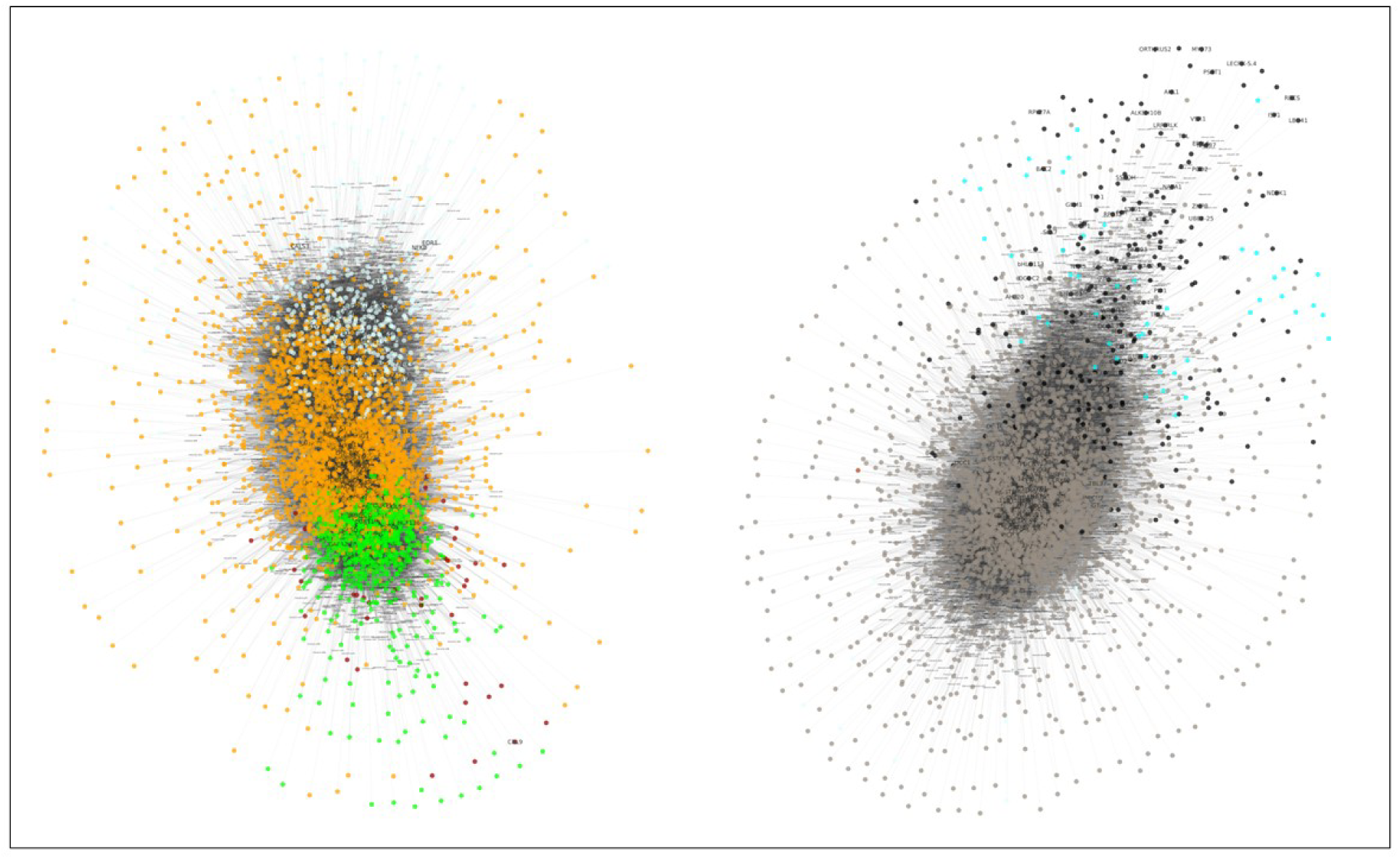
Network visualization of WGCNA modules from developmental analysis.

**Table S1.** Upregulated and downregulated genes in leaves, roots and seeds pairwise comparisons.

**Table S2.** Differentially expressed genes in leaves, roots and seeds.

**Table S3.** Differently expressed genes and their functions, arranged in clusters of expression.

**Table S4.** Hubgenes, arranged in modules of expression.

